# Spatio-Temporal Dynamics of Landscape Use by the Bumblebee *Bombus atratus* (Hymenoptera: Apidae) and its relationship with pollen provisioning

**DOI:** 10.1101/612564

**Authors:** Pablo Cavigliasso, Colin C. Phifer, Erika M. Adams, David Flaspohler, Gerardo P. Gennari, Julian A. Licata, Natacha P. Chacoff

## Abstract

Understanding how bees use resources at a landscape scale is essential for developing meaningful management plans that sustain populations and the pollination services they provide. Bumblebees are important pollinators for many wild and cultivated plants, and have experienced steep population declines worldwide. Bee foraging behavior can be influenced by resource availability and the bee’s lifecycle stage. To better understand these relationships, we studied the habitat selection of *Bombus atratus* by tracking 17 queen bumblebees with radio telemetry in blueberry fields in Entre Ríos province, Argentina. To evaluate land use and floral resources used by bumblebees, we tracked bees before and after nest establishment and estimated home ranges using minimum convex polygons and kernel density methods. We also classify the pollen of their body to determine which botanical resources they use from the floral species available. We characterized land use for each bee as the relative proportion of GPS points inside of each land use. Bumblebees differed markedly in their movement behavior in relation to nest establishment. They moved over larger areas and mostly within blueberry fields before to nest establishment, in contrast to after establishing the nest that they preferred the edges near forest plantations and changed the nutritional resources by wild floral species. Our study is the first to track queen bumblebee movements in an agricultural setting and relate movement change across time and space with pollen resource availability. This study provides insight into the way bumblebee queens use different habitat elements at crucial periods in their lifecycle, showing the importance of mass flowering crops like blueberry in the first stages of queen’s lifecycle and how diversified landscapes help support bee populations as their needs changes during different phases of their lifecycle.

## Introduction

Animal assisted pollination is crucial for the reproduction of wild and domesticated plants, and worldwide, insects are the main provider of this service [1]. Insect pollinators help to maintain trophic networks in nature [2] and help improve both quality and quantity of crops for human consumption [3–5]. Approximately 35% of global food production, and approximately 70% of economically important crop species depend upon insect pollination (to different degrees) [6–7]. Bees are one of the most important insect pollinators, but both wild and managed bee populations are declining [8–12], decreasing their potential pollination service [13–15]. Land use intensification and fragmentation associated with agriculture have contributed to bee population declines [16–17]. Understanding how bees use the resources in agricultural landscapes is essential to develop meaningful farm-based land use management plans that sustain bee populations and maximize the potential pollination service they provide to farmers and ecosystems [18–20].

In these agricultural landscapes, bumblebees (*Bombus spp*.) are one of the most important groups of bee pollinators [21]. Even so, among insect pollinators, bumblebees have experienced some of the steepest population declines and range contractions [22–25]. *Bombus spp*. have a large foraging capacity and can fly in a wider range of ambient temperatures than many other bee species [26–27], present the characteristic “buzz-pollination” causes large amount of pollen to be released, making them efficient pollinators for a variety of crops (eg. blueberry) [28–33]. They have eusocial habits [34] with colonies that can reach up to 400 individuals with several queens [35]. These have an annual lifecycle and, unlike honeybees, they do not store large quantities of honey or pollen in their nest [36]. As such, the survival of the colony depends upon the availability of suitable food for the different stages of its life cycle within foraging distance of the nest, since their nutritional requirements differ pre- and post-establishment. Environmental or habitat changes can negatively impact a colony’s success and chance of survival [37]. The forces that shape individual bumblebee flower or patch choice remain poorly understood [38], but previous work suggests that *Bombus spp*. are guided by visual, olfactory and social cues as well as the quality and quantity of floral resources [37]. This last factor resources are subject to temporal and spatial changes, presenting marked differences with respect to the stage of the cycle where they are found and translating into changes in their availability within the landscape [34]. Historically, is has been difficult to track individual bee movements [39] but today, newer technologies have enabled biologists to use miniaturized radio telemetry transmitters on bees [40–41].

We studied habitat selection of one bumblebee species, *B. atratus*, using radio telemetry in an agroecosystem dominated by blueberries in the state of Entre Ríos, Argentina. Our objective was to determine how the queens of *B. atratus* modify their spatio-temporal use of the blueberry agroecosystem [42], and to provide new knowledge about how they change their flight behavior and landscape use during different lifecycle stages. We hypothesized that the *B. atratus* queens would use landscape resources differently, changing their foraging behavior (size and shape of the home range) and the preference for certain floral resources according to the pre- and post-nesting condition. To our knowledge, this is the first study of its kind to link spatial habitat selection of bees revealed by radio telemetry with floral pollen resources in a working agroecosystem landscape.

## Materials and Methods

### Study area

The study was carried out on large-scale commercial blueberry farms in Yuqueri station, Entre Ríos province, Argentina (31°22’22.4538” S / 58°07’23.7864” W) neighboring the National Institute of Agricultural Technology, Concordia Experimental Station. The agroecosystem is characterized by the presence of blueberry and citrus fields, and small-scale *eucalyptus and pine* plantations and windbreaks.

This agro-forestry system is common and expanding in this region of northern Argentina. We conducted our study from the last week of July to the third of September 2015 when the blueberry bushes (*Vaccinium corymbosum* var. Emerald) are in peak bloom.

### Bee capture and tracking

We opportunistically netted 24 *Bombus atratus* queens that were visiting blueberry bushes at the beginning of August and September. Of the total catches, 7 queens were tagged before nesting and 10 after. We then gently immobilized the bees and glued 0.2 g radio transmitter (ATS Series A2412) on their abdomen (Fig 1.A) (S1 Text). The transmitter emits short radio pulses, allowing for real-time tracking on the ground by technicians using ATS receivers and yagi directional antennas (2.5 kHz, Advanced Telemetry Systems, Inc. R410 Reference User Manual - R06-11) (Fig 1.B). We tracked the bumblebees through the agroecosystem daily from 8 am – 6 pm for 1 – 9 days (S1 Table). Once an individual bumblebee was relocated we recorded the GPS location.

**Fig 1.**
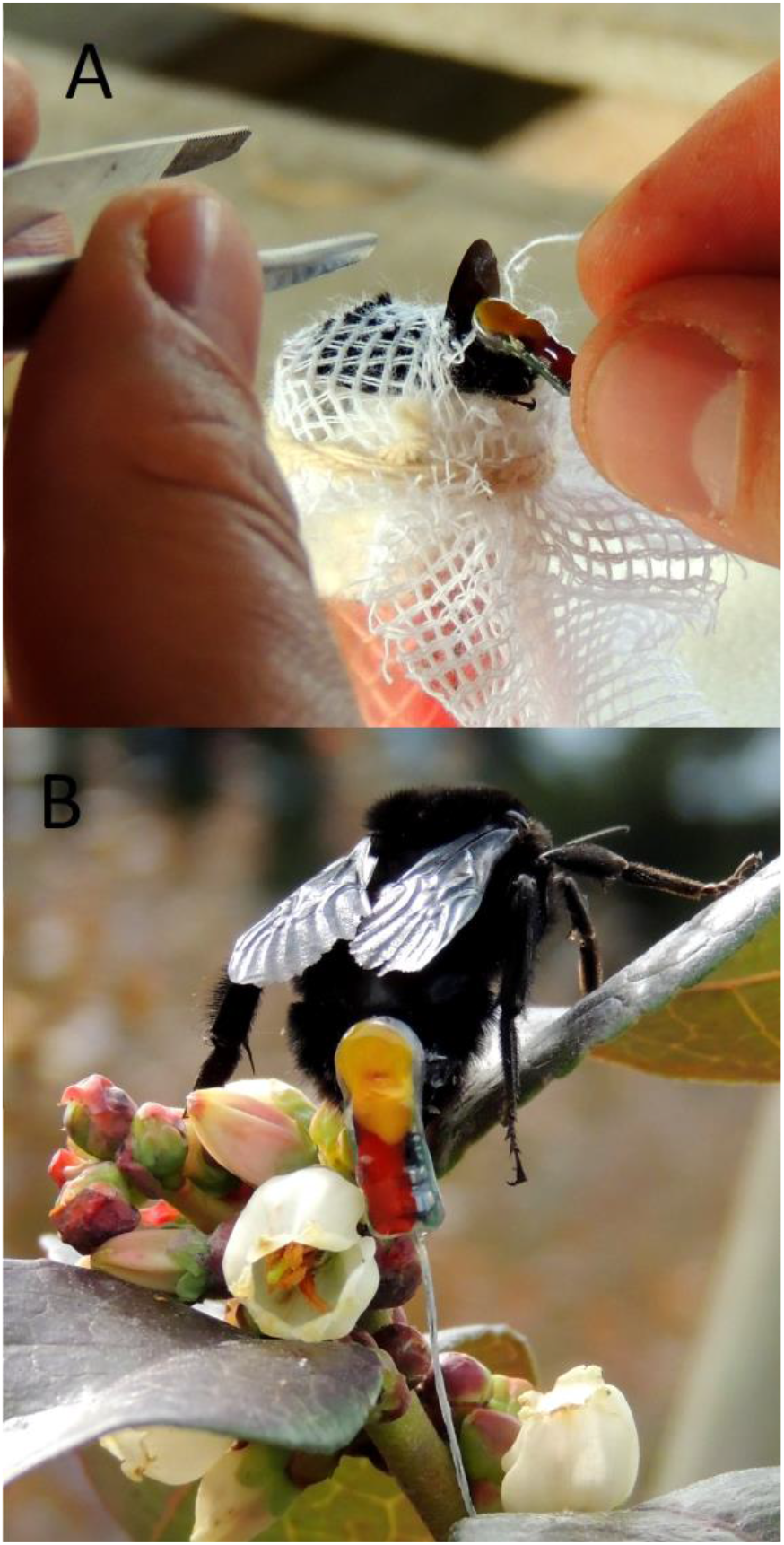
Fixing of the radius transmitter. A) Immobilization of the individual to be tracked in a soft rubber tube with a foam plunger; transmitter was attached with fast-acting glue. B) *Bombus atratus* queen with transmitter foraging on blueberry flowers. Photo credits: P. Cavigliasso.

This procedure was carried out in two different time periods of the bees’ life cycle: 1) during the nest searching location that immediately follows emergence from hibernation when the queens seek suitable a site to rear a colony; and 2) after nest establishment, when the queen has established its nest and is rearing the first cohort of workers. The nest searching period coincided with the beginning of the blueberry crop’s flowering (July 28 to August 7). The post-nest establishment period occurred during the end of the blueberry bloom and the beginning of the blooming of most native plants (August 31 to September 22) (Abrahamovich, personal observation). When a nest location was confirmed, we also recorded that location and notes its substrate.

### Land use classification

We classified the study area vegetation that cover 3,141.5 km^2^ using five land uses categories (LUC hereafter). The LUCs were grouped into: 1) *Blueberry*, the area occupied by blueberry field; 2) *Forest plantations*, comprised of planted blocks of *Pinus* and *Eucalyptus* spp. and windbreak of *Casuarina spp*.; 3) *Semi-natural area*, including pastures, abandoned lots, areas in recovery and road margins; 4) *Other fruits*, primarily citrus; and 5) *Developed*, representing human-constructions such as houses, barns and roads. The classification was done using the “Google Satellite” option of the “OpenLayers plugin” tool of QGIS (version Essen 2.14.3, available at https://www.qgis.org/es/site/), with a WGS / Pseudo Mercator projection (EPSG: 3857). We then calculated the proportional use of each land cover type based on the observed GPS locations, giving each observed point a class (e.g., blueberry or semi-natural) and quantifying the relative frequency of occurrence for each bee individual, allowing us to compare habitat use before and after nesting. These LUCs were then used in further analysis (described below).

### Bee home ranges and habitat selection

To estimate the home range and habitat selection of the queen bumblebees, we used two methods: Minimum Convex Polygon (MCP) and kernel density (KD). These two methods show complementary information on home range and habitat use, with MCPs representing the furthest ranging territory of the bees and the KD demonstrating which habitats the bees were most likely to use [43–45]. These metrics thus show us where the queens can fly and what LUC they use more intensely and thus prioritize [46].

MCP were calculated from the connected perimeter of the 5 most external recorded GPS locations taken for each individual. This method generates a polygon with an area equivalent to the minimum portion of the landscape used by each individual. From the MCP, we made inferences on the way they move, maximum flight distances, and preferences for any land use present within the landscape (land uses categories, described below). As the maximum flight distance, for each individual we used the most distant two vertices of the MCP [41]. We also characterized the shape of the polygon using two parameters: Coefficient of Compactness (Kc) and Circularity Ratio (Rci). Kc is defined as the relationship between the perimeter of a polygon and the perimeter of an area circumference equivalent to that of the polygon to be evaluated (Formula 1A – S2 Text), and is a continuous variable between 1 and 3; high values indicate very elongated areas and low values indicate more circular areas. Rci is the quotient between the areas of the polygon and that of a circle whose circumference is equivalent to its perimeter (Formula 1.B – S2 Text, range from 0-1 with 1 being totally circular areas for the unit value, square for the value 0.785 and irregular and elongated for values lower than 0.20). This coefficient is used in a complementary way for the interpretation of Kc since they describe similar parameters. These geometric parameters are widely used to classify the two-dimensional areas on maps [47–49]. These indices, although not previously used to characterize movement in animals, can be easily calculated and provide an accurate approximation of the non-uniform two-dimensional movement areas.

We calculated the KDs for *B. atratus* queens for both time periods. For this, we used the “Heatmap plugin” tool of QGIS, to create a raster layer through the density of points observed in each stage studied. For this calculation, we use the kernel function “*Quartic(triponderated)*” that resembles a circular kernel with a fixed radius to 60 layer units, which defines the direct distance from the estimated point and specifies the influence of the kernel [50]. It has been shown that this procedure is suitable for this purpose [51]. The estimators of the Kernel functions calculated for both stages are presented in S2 Table.

### Use of the floral resource around the agroecosystem

To evaluate changes in the use of floral resources before and after nest establishment, we collected queen bees each week to analyze pollen loads on their bodies, and we collected pollen from all available flowering plants in the landscape to make a pollen reference library. *B. atratus* bees were captured using an entomological vacuum while walking a random transect for 10 min in the same fields where we tracked the bees. Collected bumblebees were stored individually in Falcon tubes with 10 ml of 70% alcohol. We then collected the pollen that was adhered to bumblebee bodies by gently agitating the tube, resulting in a homogenized solution of pollen. From this solution, we extracted 10 μl, stained the pollen with Alexander’s stain, and used a Neubauer’s chamber to count the relative abundance and identity of the first 100 pollen grains observed under an optical microscope (Boeco BM-300/I/SP). Pollen found on the bumblebees was compared in three time periods following blueberry flowering and the date of capture: Early flower (4^th^ week of July and 1^st^ week of August); Peak flowering (2^nd^ and 3^rd^ weeks of August); and Post-peak (4^th^ week of August to 2^nd^ of September). The pollen library floral specimens were collected from blooming plants in the study area. Pollen samples were dried in an oven for 4 hours at 65 ° C to and we took a microphotograph of each pollen species (adaptation from Gui et al. 2014 [52]).

### Data Analysis

First, we compared foraging metrics within the condition (before and after) of nest establishment. We considered as responses variable the MCP area, maximum flight distances and shape parameters (Kc and Rci) and used a Kruskall-Wallis test.

The relative frequency of waypoints observed in each LUC during the pre- and post-nest life stages we compared through generalized linear mixed models (GLMM). For this analysis, the number of waypoints present in each LUC within the Minimum Convex Polygon (MCP) was a response variable (negative binomial distribution) and the stage (before and after establishing a nest) was a fixed effect. The analyzes were done with the statistical software R 3.5.1 (R Development Core Team, 2013). We used the *glmer* and *glmer.nb* function of the “lme4” package version 1.1-12 for the GLMM.

Finally, the number of plant species and the proportion of the pollen species best represented as indicated by pollen on bees (response variable) on every *B. atratus* queen for the three blueberry flowering time stage was compared to explore how bumblebee queens use floral resources over time. Because of the non-normal nature of these data, were completed the pollen analysis using Kruskal-Wallis test.

## Results

In total, during both study periods, we captured and tracked 24 bumblebee queens. We recorded 458 waypoints, of which 152 were obtained before bees establishing the nest and 306 were post-establishment. From bumblebees observed at the beginning of the bloom, 23 ± 11 location were recorded. In contrast with those the end of bloom that added 33 ± 20 location per queen. Seventeen bumblebees were regularly relocated (more than 5 GPS locations) and only these individuals were used for data analysis, per the criteria of the MCP and KD method.

### Foraging metrics

*Bombus atratus* changed their foraging behavior before and after nest establishment. Before selecting a nest, queens foraged over larger areas based upon MCPs (84% larger before vs. after. *H* = 6.94, *p* = 0.0068) (Table 1), with a tendency to forage within an oval shape (*H* = 1.87, *p* = 0.0702), whereas after setting a nest bumblebees queens foraged in smaller and more elongated areas. The average maximum flight distance was 642.58 ± 396.89 m, not finding significant differences between stages (*H* = 2.44, *p* = 0.1331) (Fig 2).

**Fig 2.**
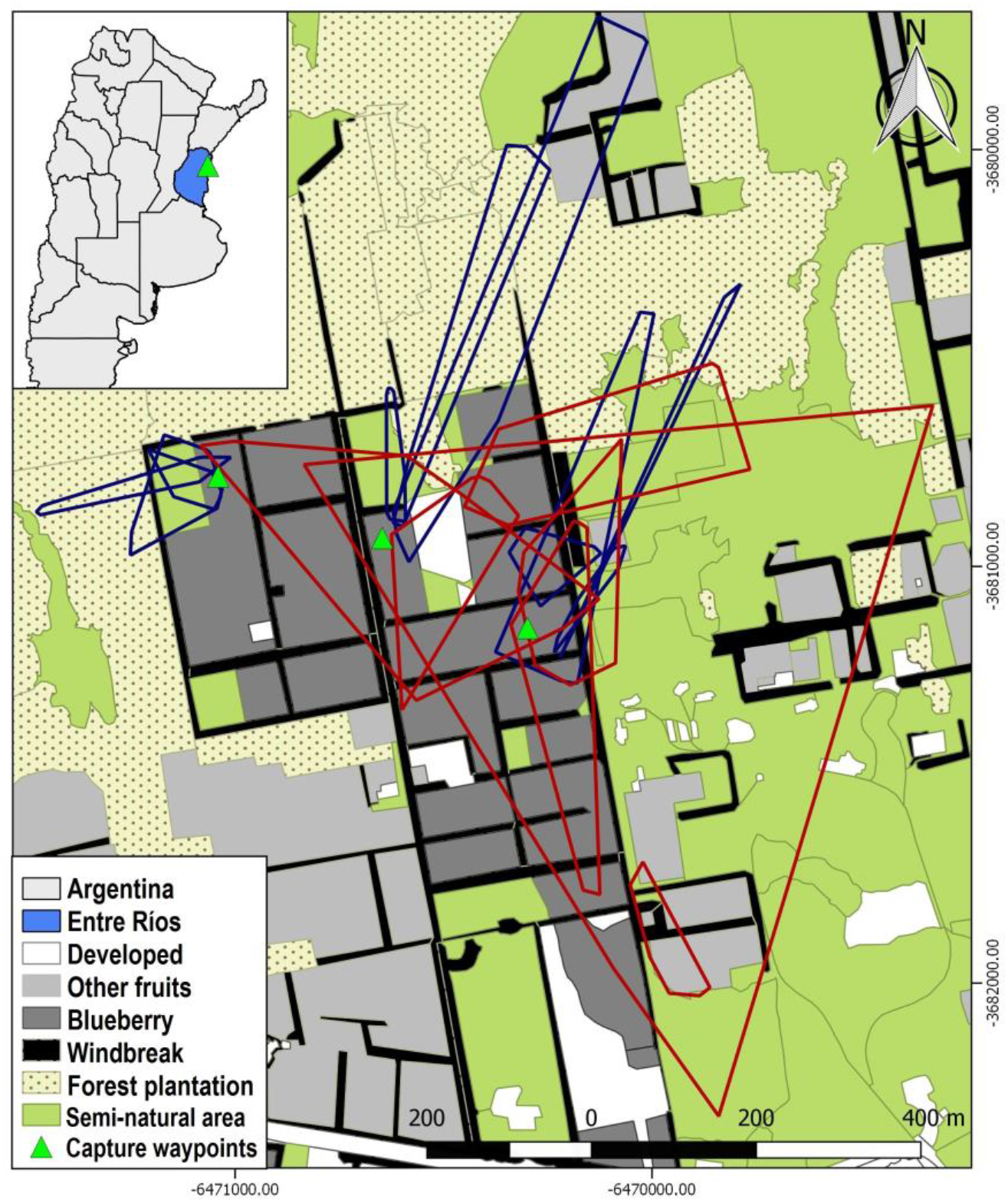
Location of the MCPs observed in both monitoring stages. Foraging areas of the queens of *B. atratus* are highlighted before (red) and after (blue) the establishment of the nests.

**Table 1.** Parameters of size and form of MCP in both stages of the home ranges of radio tracked *B. atratus* queens.

| Establishment Nest | Before | After |
| --- | --- | --- |
| MCP-Area (ha) * | 22.7 (± 31.69) | 3.56 (± 4.20) |
| MCP-Kc <sup>a</sup> | 1.35 (± 0.18) | 1.99 (± 1.08) |
| MCP-Rci <sup>b</sup> | 0.56 (± 0.13) | 0.37 (± 0.21) |
| Maximum flight distance (m) | 813.82 (± 421.77) | 522.71 (± 350.25) |

Comparison by non-parametric variance analysis Kruskal-Wallis. Mean values ( ± Standard Deviation). “\*” parameters show significant differences ( $p < 0.05$ ).
<sup>a</sup> Coefficient of Compactness

### Use of the landscape and floral resource around the agroecosystem

Before selecting a nest, queen bees focused on blueberry fields that were just beginning to flower. After nest establishment, queens tended to forage in the periphery of the blueberry, often near *semi-natural* habitats and *other fruit* LUC with blooming wild and domesticated plants (i.e., citrus plantations) (Fig 3). After nest establishment, queen bumblebees’ home ranges appear to shrink.

**Fig 3.**
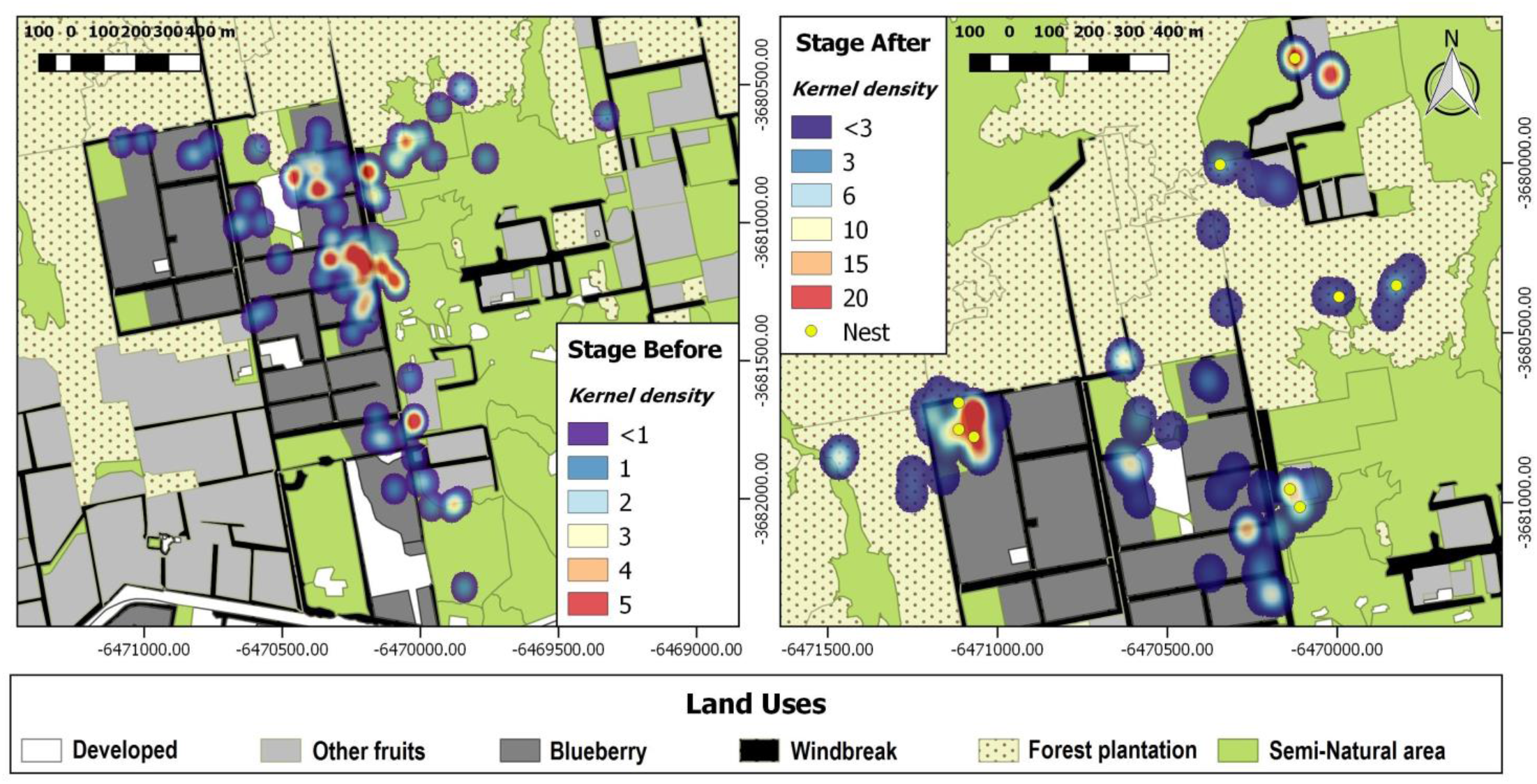
Kernel density maps of tracking bumblebees’ queens before and after setting a nest. Red values (warm colors) indicate high probability presence while Cool colors (Blue) tend to low probability values

The proportional use of different habitats differed in accordance with setting a nest. For instance, they increased their foraging in forested areas once they established a nest (GLMM. Negative Binomial. *F* = 6.11, *p* = 0.0259). Bees increased by nearly 66% their use of forest plantations (11.86 ± 4.00 %) once they have a nest (Fig 4). It should be noted that 56% of the nests observed were located on the edge (~ 3-5 m) of *Eucalyptus grandis* plantation or forest windbreaks of *Casuarina sp*., both of which are part of the plantation LUC (S3 Table).

**Fig 4.**
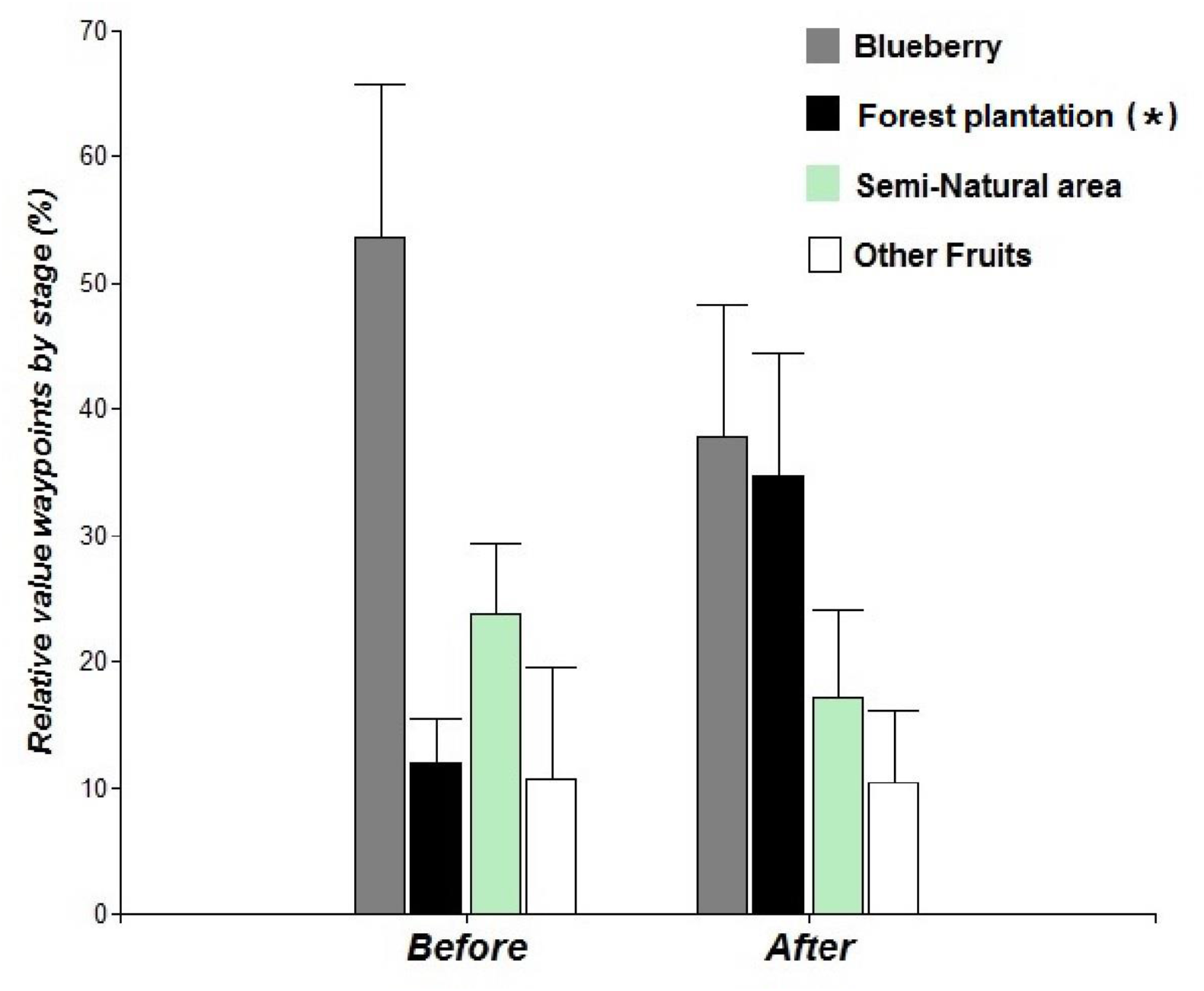
Comparison of the different LUC through GLMM. Reference with (*) = P < 0.05.

The pollen present on *B. atratus* queens (*n* = 44) captured inside the blueberry fields during the whole flowering of the var. Emerald, was from 66 plant species and did differ marginally across time of the blueberry flowering (*H* = 5.06, *p* = 0.076). During the peak of flowering of blueberry fields, the bumblebees focus their foraging on this mass flowering resource, but by the end of the blueberry flowering, other floral species increase their importance as resources for them. Plant species *Conium maculatum* L., *Buddleja stachyoides* Cham. & Schltdl. and *Nothoscordum arenarium* become more important and are collected more by queens of *B. atratus* in the post-peak period. These analyzes also show an increase in the botanical diversity of pollen present on *B. atratus* of ~ 30% more species between the peak of flowering and the post-peak (Table 2) (S4 Table).

**Table 2.**
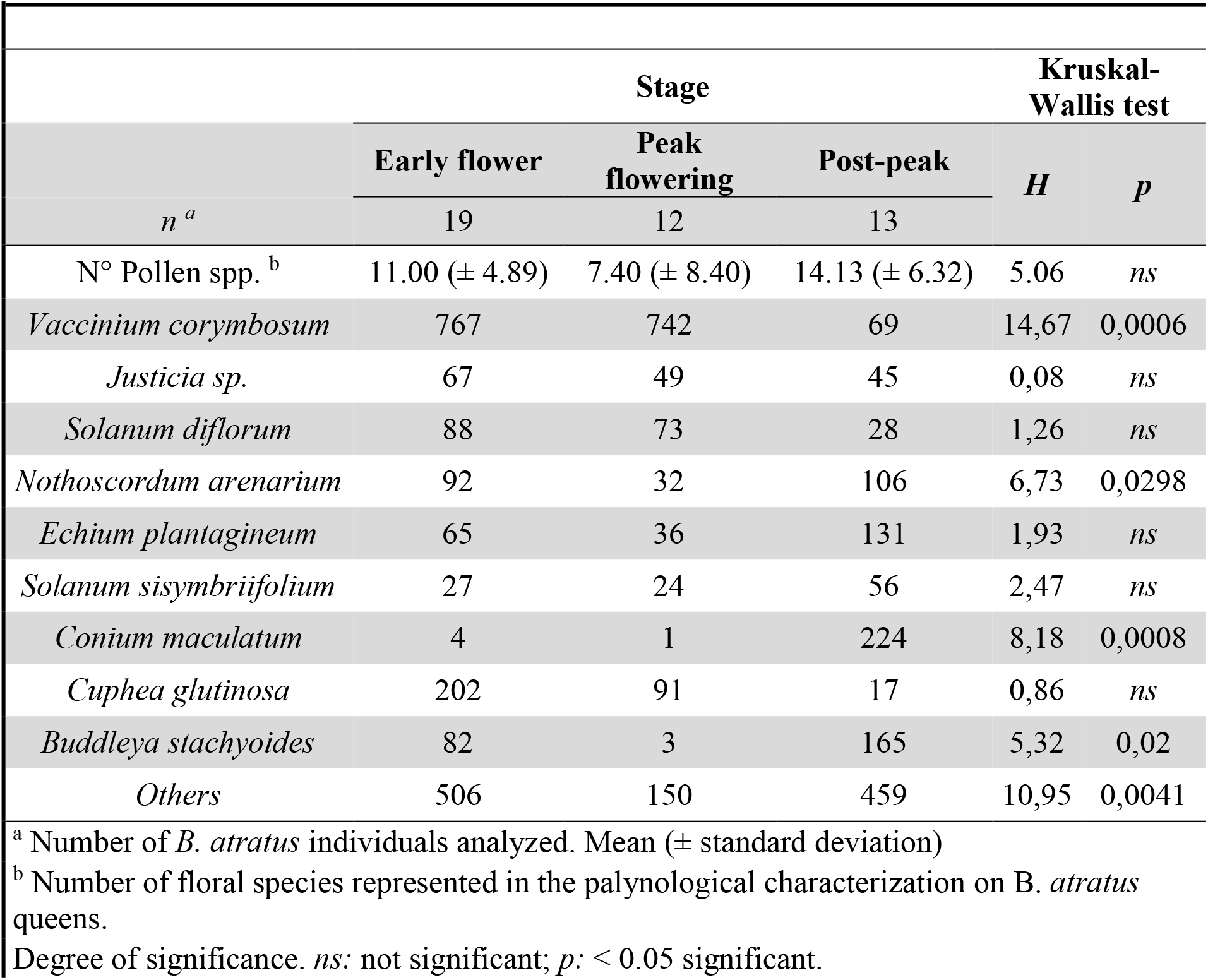
Pollen diversity and proportion of the pollen content of the most represented species on *B. atratus* at each time of flowering.

## Discussion

We investigated bumblebee habitat selection, flight distance, and home range to better understand how *B. atratus* selects floral resources in a complex and intensively used agricultural landscape. In realtime, we observed variation in the size and shape of their forage areas, flight distances, and habitat preferences related to food and nesting. Queen *B. atratus* appear to decrease their foraging areas and flight distances once they establish nests, using mostly the edges of the forest plantations to establish their colonies. During this stage, they prefer land uses with greater floral diversity to supply their growing worker colony (e.g. *Semi-Natural*). Overall, our results show the importance of a diversified habitat within agricultural areas to help sustain bumblebee’s colonies that provide pollination service to both blueberry and native plants within this region.

These results suggest two different patterns of movement for queen bumblebees during different periods in their life cycle. During the pre-nesting period, queen bumblebees flew within relatively large and circular-oval home ranges. During this life stage, queen bees often conduct reconnaissance flights of the environment in search of suitable nesting sites [27–53]. This period coincided with the beginning of the blueberry flowering, and this massive bloom likely serves as an important source of energetic resources that sustains what are likely energetically expensive nest-searching flights. Relative to some bees, bumblebees have only a modest ability to excavate a nest cavity [33]. For this reason, features correlated with variation in soil density and accumulation of leaf litter such as hedgerows, fence lines and forest edges have been found to have higher densities of bumblebee nests compared to such features as closed woods or grassland [54]. Here, we found that queens selected nest sites in habitats with a greater amount of leaf litter accumulated on the soil (i.e. windbreak and edges of plantations of *Eucalyptus* sp. and *Pinus sp*.), selecting sites adjacent to land uses with a diversity of suitable food sources and within their range of flight [55]

After *B. atratus* queens had established their nests, they changed their flight patterns and the Minimum Convex Polygons grew to be more elongated areas. In this later period, the flight behavior was more likely to be oriented with the predominant winds of spring (NW and SW), and in our landscapes this period coincides with the end of the blueberry bloom and the beginning of the other flowering plants. In these cases, the foraging sites were against the direction of the wind, suggesting that the bees had established sources of pollen resources. This behavior has been documented for individual bees who know their environment and have standardized routes of foraging (i.e. “bee line”) [56–57]. At this later stage, the queens were likely focused on collecting pollen in mass to feed the growing worker bee population that would soon emerge.

In the same way that the requirements of the species of floral visitors are modified during their life cycle, the supply of nutritional resources that the environment provides generally changes, forcing to have an adaptive behavior by of species that collect their food from renewable resources [58]. This study is a snapshot in time of how *B. atratus* queens modified their interactions with the habitat before and after the formation of nest. During the nest searching period the queens intensely used the blueberry fields since the flowers provide rich and abundant nectar and pollen. Following nest establishment, queens care for their emerging worker bees and are reducing their movements [33]. At this stage of their life cycle, the nutritional requirements for the queen and the colony change. The future worker bees require protein-rich food for its development [59]. Consequently, the bees’ movements shifted to include the land use categories with greater pollen heterogeneity [60–61] despite continued, albeit reduced, availability of blueberry flowers.

*Bombus atratus* movements are similar to those reported for other bumblebees from Europe (S5 Table). Few studies have studied the flight behavior in *Bombus* queens, finding results similar to those obtained by Walther-Hellwig & Frankl (2000) [62] by the capture-recapture method for *B. terrestris* and ~ 50% less than those observed by Hagen et al. (2011) [41] using telemetry technology in queens of *B. hortorum*. Likewise, more studies of movements in this bumblebee caste are lacking to be able to specify a flight pattern and generalized foraging behavior for these stages of its life cycle.

The results obtained from the queens of *B. atratus* around the blueberry agroecosystem demonstrate how they change the size and shape of their home ranges, but also the use of land use categories as their dietary needs change. Although the relative presence of bumblebees in land use groups in general does not show significant differences, after the establishment of a nest, forest plantations emerge as an important habitat feature, increasing their use by more than 65% and housing 56% of nests observed. This observation suggests that these small-scale plantations can represent a valuable resource for this species providing shelter and possible nutrients [63–64]. The plantations may also serve as guides in foraging flights since bumblebees are more likely to perform straight flights when flying along windbreak compared to when they are flying in open fields, suggesting that they may follow linear landscape features [65]. In addition, these actors are actively pollinating within the fields at a time when there are not many other species of native pollinators, giving them an intrinsic value in this agroecosystem

The analysis of body pollen reinforces our telemetry experiment, showing that between the periods of blueberry bloom there was a variation in the pollen proportion of floral species collected from the bumblebees. In the post-peak blueberry period there was an increase of 30+ % in the diversity of pollinic morphotypes present on the bumblebees. This result suggests that they looked for food in the other land use categories to meet the changing nutritional needs of the workers. It should be noted that, the Emerald variety of blueberry planted in the fields is the first to bloom in the region and conventional blueberry production systems may combine batches of different varieties with subsequent or sequential flowering curves. This observation supports our hypothesis that the *B. atratus* queens change how they use the available landscapes based upon the resource availability and perform a cost-benefit evaluation according to the nutritional needs required by the stage of their life cycle [66–70]. This is likely one of the most sensitive stages of the bumblebee’s life cycle, aggravated when there is a shortage of resources for foraging, which could cause the death of the young queen and her colony [34]. In this context, the massive bloom of blueberry fields emerges as an important source of nectar and pollen in this period, supporting the establishment of new colonies.

### Final considerations

This is among the first studies to link flight behavior with floral and nesting resources in a productive mosaic agroecosystem, and demonstrates how the resource needs of bumblebee queens’ changes over time and relies on semi-natural areas surrounding agricultural fields as foraging habitat. Heterogeneous landscapes can provide diverse resources that are needed by *B. atratus* queens at different moments of their life cycle. Blueberry fields appear to be an important resource at the beginning of their life cycle until the moment of nesting. At the same time, the edges of forest plantations seem to offer nesting habitat for native bees when they are adjacent to pollen-rich fields, and the semi-natural areas are harnessed for the workers’ protein-rich diet. We emphasize that we did not directly observe the bees using the bare soil or the land uses developed during our study.

Bees provide vital ecosystem services as pollinators and we need to work to sustain these wild pollinators. The management and conservation of these semi-natural land use categories is an important part of achieving sustainability of agro-ecological systems because they help supplement bee nutritional needs with diverse pollen sources [71] and nesting sites. Semi-natural habitats provide essential resources for the formation and survival of the worker caste that, when upon emerging, will take the lead in supplying the colony with pollen, and thus providing for the next season’s queens [72]. Our work contributes to the growing understanding of how bumblebees use the environment, and provides valuable information for conservation planning and sustainable management of the land at a crucial moment in its life cycle. We suggest that land owners and managers of agricultural lands should consider the full life cycle of bees from nest formation to the worker bee emergence, and this longer-term perspective can help maintain native bees in farmlands from year after year, maximizing the pollination service they provide.

## AUTHORS CONTRIBUTIONS

P.C., C.C.P., E.A. and J.L. conceived the ideas and jointly designed the telemetry methodology while P.C. designed the pollen analysis portion of the project; P. C., C. C. P. and N. P.C. contributed equally to the writing of the manuscript; P. C., E. A. and C. C. P. collected the samples; P. C. analyzed the data. This work was supported by the National Science Foundation Partnerships in International Research and Education grant (No. 1243444) to D.F. All authors contributed critically to the drafts and gave final approval for publication.

## Acknowledgments

This research was supported by the US National Science Foundation Partnerships in International Research and Education research grant num. 1243444. We thank Fernanda Rivadeneira and all the members of the Fruit and Forestry Area of the Experimental Station Concordia for helping with field logistics and for connecting us with area landowners, and for the use of equipment and vehicles. Lastly, we thank Nicolas O. Monzon, for the assistance and good predisposition in the collection of wild flora and identification of the different floral species pollen.

